# A small interfering RNA (siRNA) database for SARS-CoV-2

**DOI:** 10.1101/2020.09.30.321596

**Authors:** Inácio Gomes Medeiros, André Salim Khayat, Beatriz Stransky, Sidney Emanuel Batista dos Santos, Paulo Pimentel de Assumpção, Jorge Estefano Santana de Souza

## Abstract

Coronavirus disease 2019 (COVID-19) rapidly transformed into a global pandemic, for which a demand for developing antivirals capable of targeting the SARS-CoV-2 RNA genome and blocking the activity of its genes has emerged. In this work, we propose a database of SARS-CoV-2 targets for siRNA approaches, aiming to speed the design process by providing a broad set of possible targets and siRNA sequences. Beyond target sequences, it also displays more than 170 features, including thermodynamic information, base context, target genes and alignment information of sequences against the human genome, and diverse SARS-CoV-2 strains, to assess whether siRNAs targets bind or not off-target sequences. This dataset is available as a set of four tables in a single spreadsheet file, each table corresponding to sequences of 18, 19, 20, and 21 nucleotides length, respectively, aiming to meet the diversity of technology and expertise among labs around the world concerning siRNAs design of varied sizes, more specifically between 18 and 21nt length. We hope that this database helps to speed the development of new target antivirals for SARS-CoV-2, contributing to more rapid and effective responses to the COVID-19 pandemic.

## INTRODUCTION

Started in late December 2019, coronavirus disease 2019 (COVID-19) rapidly transformed into a global pandemic, with an incidence of more than 30M cases and almost 1M deaths around the world as of September 2020^1^, and strongly negatively impacting the global economy (1). This circumstance brought a huge demand for developing antivirals capable of targeting the SARS-CoV-2 RNA genome and RNA interference approaches (2–4) emerged as a possible solution. Small interference RNA (siRNAs) are RNA sequences about 20nt-long that, together with RNA-Induced Silencing System (RISC) (6), bind interest mRNA molecules (4, 5) inhibiting its translation and expression.

RNAi approaches have been employed for SARS-CoV (6–8), with reports of viral levels decreasing (9), and recent works claim that it may also work for SARS-CoV-2 (10, 11). Researchers in (12) used Immune Epitope Database and Analysis Resource (IEDB) to find potential regions in diverse coronaviruses with matches to SARS-CoV-2, identifying many of them in SARS-CoV, the closest homolog. Chen *et al* (13) apply a window of 3000 nucleotides with a step of 1500 over reference SARS-COV-2 genome (MN908947^2^) seeking 1–25nt regions called “free segments”. Besides, siRNAs databases targeting a broad range of viruses (14–16) have been developed. Recently, researchers developed a SARS-CoV-2 oligonucleotide sequence database, to improve the SARS-CoV-2 detection and treatment methods, providing sequences with the lowest and highest conservation levels (17).

In this work, we propose a SARS-CoV-2 targets database to support siRNA approaches, aiming to speed up RNAi design by providing a set of possible targets and siRNA sequences with the required information for choosing the most appropriate targets for new siRNAs. Unlikely cited databases, which are manually curated, we apply a sliding-window approach for covering whole SARS-CoV-2 genomic space, extracting every possible siRNA sequence of 18, 19, 20, and 21 nucleotides, enabling researchers to assess solutions capable of targeting any region of the virus. The database has more than 170 features, including thermodynamic information, base context, target genes, and alignment information against diverse SARS-CoV-2 strains, together with scores and predictions collected from three siRNA efficiency prediction tools. All this coordinated information will enable users to select with higher confidence targets that best match a broad set of conditions for designing even more efficient siRNAs.

## MATERIAL AND METHODS

Although siRNAs length can vary from 18 to 25 nucleotides (18), synthetic ones should range from 19 to 21nt (19), according to ThermoFisher siRNA Design Guidelines^3^. Thus, the proposed database provides information about each possible 18 to 21 nucleotides siRNA target region from SARS-CoV-2, one table for each length. Moreover, tools employed for assessing siRNAs efficiency (20–22) operate over sequences lying in that range, which reinforces our choice. Since they present the same columns, we explain here the development process only for the 21-length table.

SARS-CoV-2 reference genome was collected from NCBI (code NC_045512) and a sliding window of 21nt-long and step 1^4^ were used to traverse the genome. Table 1 indicates the total number of sequences obtained for each length. Seven new sequences sets were then generated from the obtained sequences set (called *target region*), following the aforementioned ThermoFisher guidelines, and suggestions from our collaborators: (i) *natural sense*, by removing the first 5’-end dinucleotide from *target region* sequences; (ii) *oligo natural sense*, replacing thymine with uracil over *natural sense* set; (iii) *synthetic sense*, by replacing first 3’-end dinucleotide from *natural sense* sequences with TT; (iv) *oligo synthetic sense*, by replacing of thymine with uracil over *synthetic sense* sequences; (v) *antisense*, from the reverse complement of *target region*; (vi) *oligo antisense*, by replacing thymine with uracil over *antisense* sequences; and (vii) *oligo antisense rev* set, by reversing *antisense* sequences.

**Table 1.** Number of siRNAs of each length.

| Length | Number of siRNAs |
| --- | --- |
| 18 | 29,883 |
| 19 | 29,882 |
| 20 | 29,881 |
| 21 | 29,880 |
| Total | 119,526 |

*Natural sense* sequences were then aligned against (a) SARS-CoV-2 reference genome, to verify which genes they align with; (b) the human genome (NCBI accession code GRCh37) and coding and non-coding transcriptome, to verify potential cross-reaction with off-target transcripts; (c) SARS-CoV-2 strains available at GISAID^5^ initiative coming from Brazil, China (Wuhan region only and whole country less Wuhan), England, Germany, Italy, Russia, Spain and USA; and (d) reference genomes of MERS virus (NCBI MG987420), SARS-CoV (NCBI NC_004718) and Influenza virus genome (NCBI NC_026438), aiming to assess whether siRNAs are capable to target regions from those viruses and strains. Bowtie^6^ (23) version 1.1.0 was used as the aligner, using the following flags: -a, -S, --pairtries equals to 4, -p equals to 40, -n equals to 3, -l equals to 7 and -f. Flag -e was used, being equals to 150 when aligning against the human genome, 10 for GISAID strains, and 220 for remaining genomes. Sequence properties regarding base context and alignment information were calculated from the above sequences sets and performed alignments (see Supplementary Text 1). Thermodynamic information and expected efficiency of candidates siRNA designed for targeting those regions was calculated with OligoCalc^7^ (24) and three predictors, namely ThermoComposition21 (20), SSD (20, 22), and si-shRNA Selector (21),

## RESULTS

### Database analysis & statistics

The proposed database displays a total of 119,526 siRNAs divided in four different sizes ranging from 18 to 21 nucleotides (see Table 1 for the number of siRNAs of each length). As stated, we applied over them three siRNA efficiency prediction tools to assess their inhibition power. Figure 1 illustrates the number of 21nt *antisense* siRNA sequences predicted as effective by every single predictor, and the quantities predicted by more than one. It can be seen that no siRNA was unanimously considered effective, while approximately 53% of them (15,821 siRNAs) were considered as such just by SSD. Besides that, a single siRNA was predicted as effective by both ThermoComposition21 and si_shRNA_selector.

**Figure 1.**
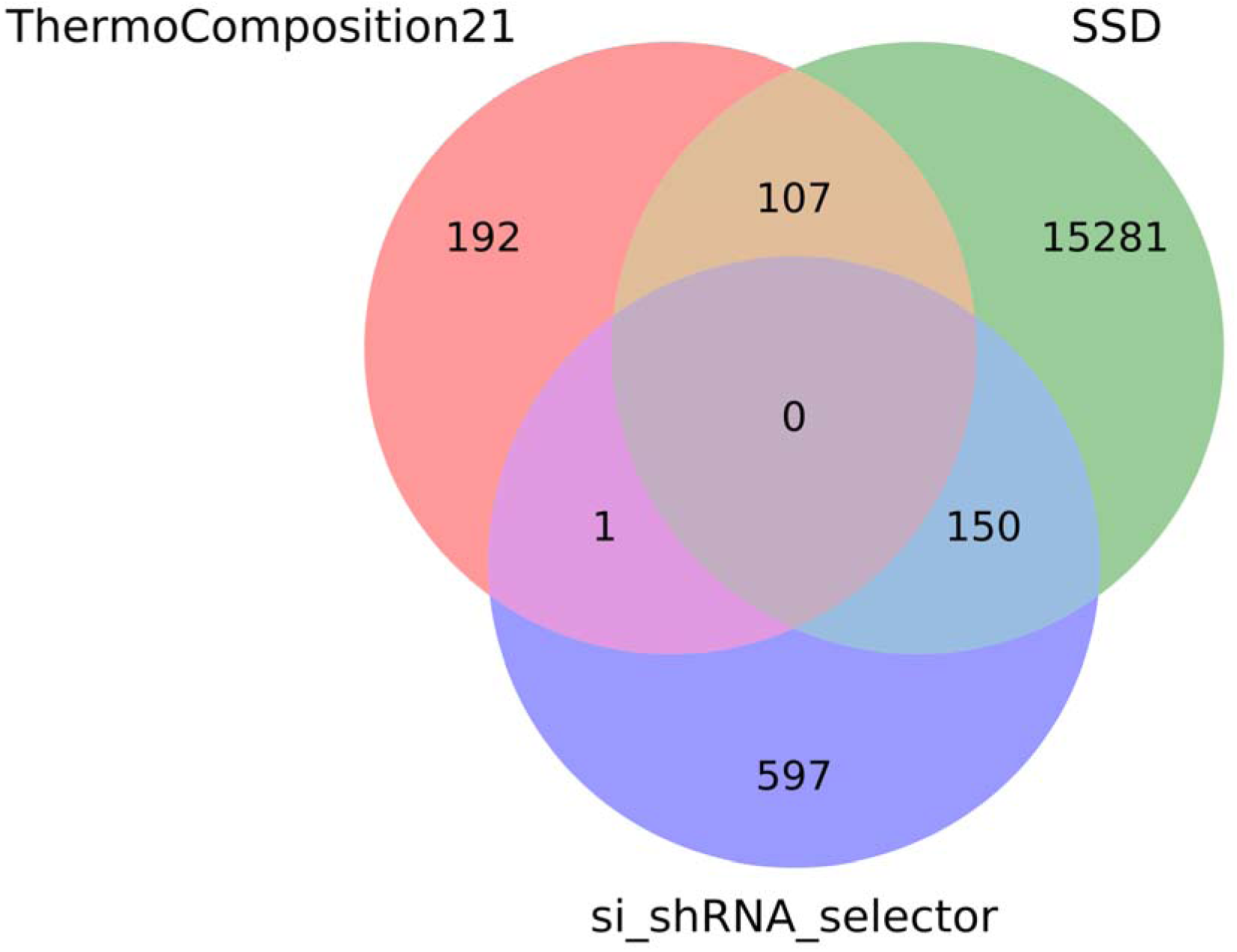
Venn diagram of three predictors for 21nt-long antisense sequences classified as efficient.

We also aligned all siRNAs to the Human genome, Human coding and non-coding transcriptomes, SARS, MERS, and H1N1 genomes, with Bowtie version 1.1.0, aiming to identify if siRNAs could off-target regions in those organisms, thus presenting cross-reactivity with them. Figures 2a-d illustrate the growth of siRNAs quantities as the minimum number of necessary mismatches to have alignment increases. The results show that virtually all 18nt siRNAs can match the human genome and transcriptomes (coding and non-coding) with three mismatches. This number, however, increases to four considering 19nt and 20nt (Figures 2b and 2c), and to five considering the 21nt length (Figure 2d). It can also be noted that, in each length, about 2,500 siRNAs match perfectly with some region of SARS. Regarding MERS and H1N1, about 2,500 18nt siRNAs can match regions of those viruses, when the minimum number of allowed mismatches is two (Figure 2a). This number is only overcome by 19-21nt siRNAs only when the number of mismatches is increased to three (Figures 2b-d). Finally, it can be observed that while all 18nt and 20nt siRNAs match some regions from MERS, SARS, and H1N1 using at least six mismatches, the number of mismatches increases to seven for 19nt and 21nt siRNAs.

**Figure 2.**
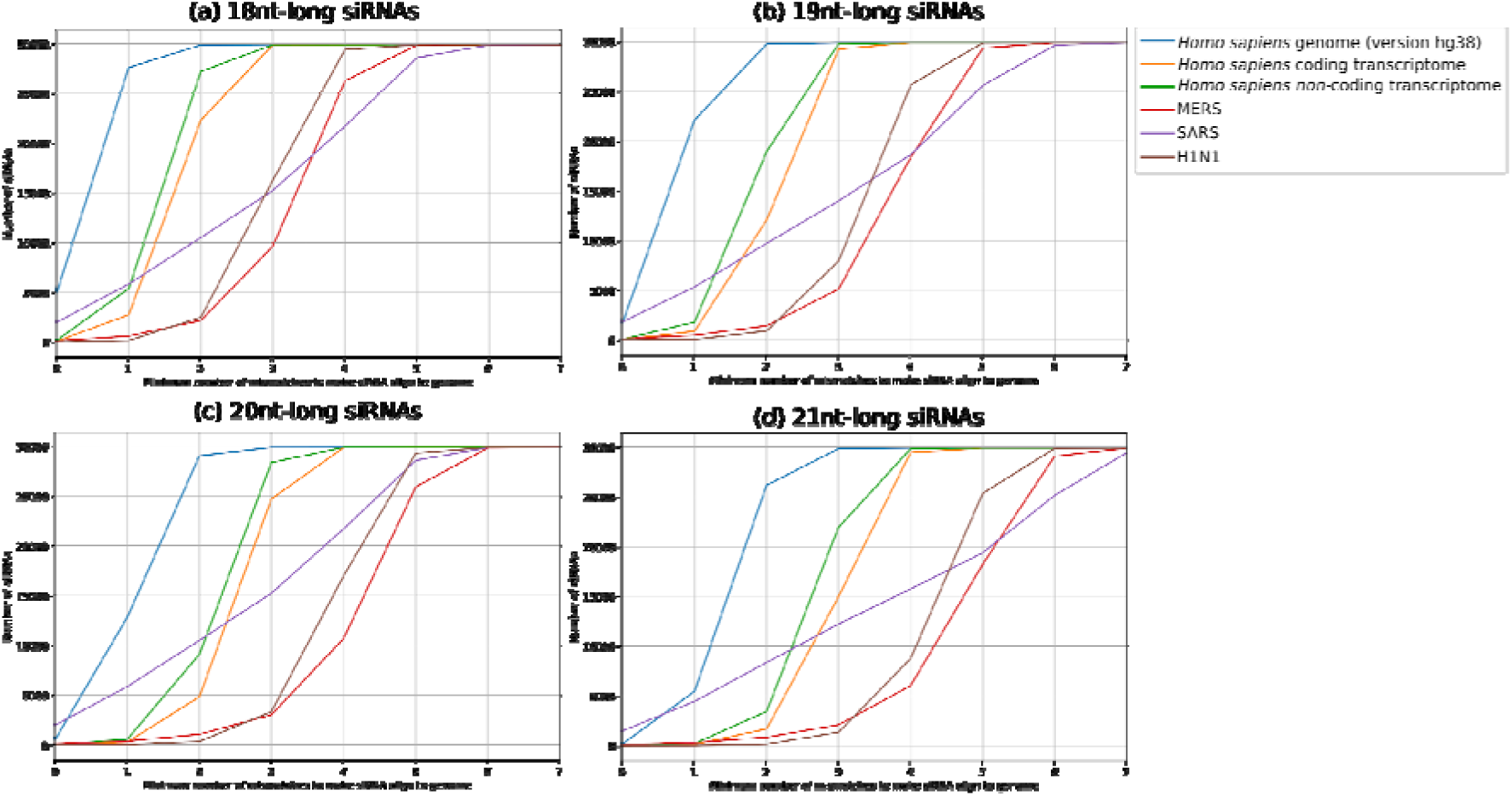
Number of antisense (a) 18nt-long, (b) 19nt-long, (c) 20nt-long, and (d) 21nt-long siRNAs with different mismatches against Human genome, Human coding and non-coding transcriptome, and MERS, SARS, and H1N1 genomes.

### Database use and access

The proposed database is distributed as a spreadsheet file containing four tabs, each one corresponding to *target region* sequences of a specific length. Here we will present how a researcher can use this database with an illustrative example. Suppose a user wants to select siRNAs with 21 nucleotides length. In this case, the user will access the tab “21 bases” from the spreadsheet file. After opening it in a spreadsheets editor, the next step is selecting siRNAs whose properties match the user requirements. Assume that the user wants a siRNA that has little or no homology with the human genome, can act over as much as possible British SARS-CoV-2 strains, and its first dinucleotide is AA. This last requirement is achievable by applying a filter over column P to show only lines with value 1 on it (see Supplementary Text 1), decreasing the number of siRNA candidates from 29880 to 2858. For little or no homology with the human genome, the number of mismatches against human sequences must be at least three (25). Filtering table to display lines with at least a value of three at columns BO, BP, BQ, CG, CH, CI, CY, CZ, and DA (Supplementary Text 1) now reduces candidates from 2858 to 999. Finally, to approach as many British strains as possible, column BY (Supplementary Text 1) can be filtered to display only the three highest values, for example, which reduces candidates from 999 to 10 candidates. Such a reduction not only saves wet-lab tests costs but also ensures that selected siRNAs meet the main user requirements.

## DISCUSSION

Designing siRNAs is a challenging procedure, because sometimes minor changes in its nucleotide sequence can alter its functionality (26). As reported in (27), specificity, potency, and efficacy of siRNA-mediated gene silencing can be determined by analyzing siRNA nucleotide sequence, hence its inability to bind to unintended regions (off-targets) is an important factor that must be strongly taken into consideration. Therefore, we proposed a SARS-CoV-2 targeted siRNAs database with sequence and thermodynamic stability information, to help the evaluation of important factors related to their efficacy and optimize the decision process towards choosing the best ones as target antiviral solutions. Considering that each laboratory has its own technology context and expertise in designing siRNAs of specific lengths, we provide a list of siRNAs varying from 18 to 21 nucleotides-length, aiming to meet the range of possible lengths used in the design process.

Numerous works have been proposing methods and guidelines for choosing the best siRNAs by analysing their sequence characteristics (28–31), for which two broad reviews are available at (26, 32). Our proposed database provides information regarding base, GC and AU context, so as the quantities of each RNA nitrogenated base in sequences, besides information about the presence of UUUU and GCCA, considered toxic motifs (33) So any user with a proper efficacy evaluation method (or anyone provided by literature) can easily evaluate siRNAs with this database at disposal. It also provides thermodynamic information collected from the application of three predictors (20–22), thus enabling users to have a deeper look at siRNAs’ properties, and choose the best ones according to their specificities. As it can be seen in Figure 1, they have high divergence when setting a siRNA as efficient or not, which suggests that they must be used in a complementary way. Due to genetic diversity and variability of SARS-CoV-2 (34), a siRNA that is highly efficient over one strain may not be when applied to another. Hence, we also provide similarity information with strains from diverse countries, such that users will benefit from the opportunity of input geographical specificity and even more customization to their decision process.

Ensuring that siRNAs are not capable of targeting human sequences (off-targets) is also another important requirement, for which a minimum of three mismatches is necessary to meet it (25). Thus, similarity information with the human genome, coding and non-coding transcriptome, is also available in our database. As it was shown in the Database Analysis & Statistics session, virtually all 18nt-long siRNAs matched with such genome and transcriptomes with at least three mismatches, corroborating aforementioned statement from literature. For the best of our knowledge, this is the first database to figure siRNAs similarity information against human coding and non-coding transcriptomes, giving to users even more confidence power about siRNAs specificity. It is hoped, with this database, that the development of new target antivirals for SARS-CoV-2 using RNAi technology can be not only eased and accelerated, but also capable of identifying even more efficient solutions for silencing that virus, and contributing to the control of the pandemic.

We made available the proposed database as a spreadsheet file given the urgency to provide this information for the scientific community that is developing effective therapeutics for SARS-CoV-2. Additionally, it is intended to build a webpage for more user-friendly and interactive access to the data. Moreover, we also intend to replicate the approach employed in this work for exploring the genomic space of other viruses, as well as ones that may represent a threat to possible new pandemic events.

## AVAILABILITY

A spreadsheet file regarding database tables is available at Open Science Framework (https://doi.org/10.17605/OSF.IO/WD9MR) and mirrored at http://www.bioinformatics-brazil.org/siRNAdb/sirnas_cov_db.xlsx. Codes and binaries regarding software employed to build the database are available at https://github.com/inaciomdrs/sirna_db_building_protocol. A protocol describing technical details about database generation is currently submitted and under consideration to Nature Protocol Exchange.

## ACKNOWLEDGEMENT

We acknowledge the Pró-Reitoria de Pesquisa from Universidade Federal do Rio Grande do Norte and Pró-Reitoria de Pesquisa from Universidade Federal do Pará. We also acknowledge the Bioinformatics Multidisciplinary Environment (BioME) at UFRN and Bioinformatics Graduate Program, IMD/UFRN for the provision of computational resources.

## Supplementary Text 1 - Features description

A set of more than 170 features was calculated for each sequence from *target region* together with their derivations in six new sets. The first column, A, provides a produced ID for the sequence, and columns B, D, E, G, H, J, K, and L, the proper sequence, and its correspondents in seven sets. Column N indicates SARS-CoV-2 genes contemplated in *target region* sequence. Columns P to S indicate, respectively, if the sequence from *target region* starts with AA, TT, GG, or CC (1 if it starts, 0 otherwise). Likewise, columns U to X indicate which of AA, TT, CC, and GG dinucleotides *target region* sequence ends (1 if it ends, 0 otherwise).

Columns AA to AI contains the following information for *oligo natural sense* sequence: columns AA to AD gives the quantities of adenine, uracil, guanine, and cytosine, respectively; columns AE and AF, GC and AU% content; columns AG and AH, if tetranucleotides UUUU and GCCA, respectively, are present in sequence (1 for present, 0 otherwise); and column AI, the number of pentamers^8^ (in the sequence) that are AU-rich (at least 80% of its composition is AU). This same set of features is also calculated for *oligo synthetic sense* sequences (columns AL to AT) and *oligo antisense* ones (columns AW to BE). Column BG provides AU% content of the first heptamer from *oligo antisense rev* sequence, while column BH gives GC% content from the last heptamer of it.

Columns BJ to BL indicates if there is in *natural sense, synthetic sense*, and *antisense* sequences, respectively, any 6-length palindromic window (1 for true, 0 otherwise). Columns BO to BT presents the minimum number of mismatches needed for *natural sense* sequence have homology with the human genome (NCBI accession code GRCh37) and coding and non-coding transcriptome, MERS genome (NCBI accession code MG987420), SARS genome (NCBI accession code NC_004718), and Influenza virus genome (NCBI accession code NC_026438); and columns BV to CD, the respective number of SARS-CoV-2 strains genomes^9^ from Brazil, China (Wuhan region only), China (without Wuhan region), England, Germany, Italy, Russia, Spain and USA that *natural sense* sequence has homology with. Columns CG to CV replicate this information set for *synthetic sense*; and CY to DN, for *antisense* sequence.

Columns DQ to DZ bring thermodynamic information provided by OligoCalc^10^ (1) to *oligo natural sense*, namely, melting temperature, melting temperature adjusted considering a Na^+^ concentration of 50mM, melting temperature calculated as described in (2) using the values available in (3), product between general gas constant *R* and natural logarithm of 1 over primer concentration, sequence ΔG (change of oligonucleotide’s free energy), sequence ΔH (change of enthalpy), sequence ΔS (change of entropy), number of potential hairpin sites, number of potential self-annealing sites, and whether sequence 3’ have self-complementarity (1 if yes, 0 otherwise). Columns EC to EL replicate this information set for *oligo synthetic sense* sequence, and columns EO to EX, for *oligo antisense* sequence.

Columns FA to FJ bring efficiency score and thermodynamic information provided by *ThermoComposition21* program (4) to *natural sense* sequence, namely, predicted effectiveness^11^, number of GG dinucleotides present in the sequence, and a set of eight thermodynamic indexes (measured in ΔG) calculated by the tool which is used for effectiveness prediction. Columns FM to FV replicate this information set for *synthetic sense* sequence, and columns FY to GH, for *antisense* sequence.

Column GJ provides predicted efficiency^12^ from SSD program (5) over *natural sense* sequence, and columns GL to GO, a set of four thermodynamic indexes (measured in ΔG) calculated by the tool for *natural sense* sequence, which are used for effectiveness prediction. Columns GR to GU replicate this information set for *synthetic sense* sequence, and columns GX to HA, for *antisense* sequence.

Finally, columns HD to HF bring efficiency prediction and thermodynamic information provided by software *si-shRNA Selector* program (6) to *natural sense* sequence, namely, predicted efficiency^13^ and a set of two thermodynamic indexes (measured in ΔG) calculated by the tool which are used for effectiveness prediction. Columns HI to HK replicate this information set for *synthetic sense* sequence, and columns HN to HP, for *antisense* sequence.

## Footnotes

1 https://www.who.int/docs/default-source/coronaviruse/situation-reports/20200805-covid-19-sitrep-198.pdf?sfvrsn=f99d1754_2

2 https://www.ncbi.nlm.nih.gov/nuccore/MN908947

3 https://www.thermofisher.com/br/en/home/references/ambion-tech-support/rnai-sirna/general-articles/-sirna-design-guidelines.html

4 This parameter is used in all tables, independently of length

5 https://www.gisaid.org/

6 http://bowtie-bio.sourceforge.net/

7 http://biotools.nubic.northwestern.edu/OligoCalc.html

8 Obtained through a traversing over sequence with a 5-long one step sliding window.

9 The header of each column displays the total number of strains from each country that sequences were aligned to.

10 In order to apply it to a huge volume of sequences, we have translated javascript code of webserver related to thermodynamic information calculus to in-house Python (https://www.python.org) scripts.

11 Quantified by its gene silenced activity, ranging from 0 (complete gene knockout) to 100 (no effect).

12 Categorical field, where 1 means that sequence in question is an efficient siRNA, 0 otherwise.

13 Categorical field, where 1 means that sequence in question is an efficient siRNA, 0 otherwise.

